# The psychological correlates of distinct neural states occurring during wakeful rest

**DOI:** 10.1101/2019.12.21.885772

**Authors:** Theodoros Karapanagiotidis, Diego Vidaurre, Andrew J. Quinn, Deniz Vatansever, Giulia L. Poerio, Adam Turnbull, Nerissa Siu Ping Ho, Robert Leech, Boris C. Bernhardt, Elizabeth Jefferies, Daniel S. Margulies, Thomas E. Nichols, Mark W. Woolrich, Jonathan Smallwood

## Abstract

When unoccupied by an explicit external task, humans engage in a wide range of different types of self-generated thinking. These are often unrelated to the immediate environment and have unique psychological features. Although contemporary perspectives on ongoing thought recognise the heterogeneity of these self-generated states, we lack both a clear understanding of how to classify the specific states, and how they can be mapped empirically. In the current study, we capitalise on advances in machine learning that allow continuous neural data to be divided into a set of distinct temporally re-occurring patterns, or states. We applied this technique to a large set of resting state data in which we also acquired retrospective descriptions of the participants’ experiences during the scan. We found that two of the identified states were predictive of patterns of thinking at rest. One state highlighted a pattern of neural activity commonly seen during demanding tasks, and the time individuals spent in this state was associated with descriptions of experience focused on problem solving in the future. A second state was associated with patterns of activity that are commonly seen under less demanding conditions, and the time spent in it was linked to reports of negative rumination. Finally, we found that these two neural states tended to fall at either end of a neural hierarchy that is thought to reflect the brain’s response to cognitive demands. Together, these results demonstrate that approaches which take advantage of time-varying changes in neural function can play an important role in understanding the repertoire of self-generated states. Moreover, they establish that important features of self-generated ongoing experience are related to variation along a similar vein to those seen when the brain responds to cognitive task demands.

## Introduction

When unoccupied by an external task, humans often engage in complex patterns of self-generated thinking. Often studied under the rubric of mind-wandering (Smallwood and Schooler, 2006, 2015) or daydreaming (Singer, 1966), these patterns of thought often share a common feature of being unrelated to events in the here and now. However, they are also heterogeneous in content (Smallwood and Andrews-Hanna, 2013) and outcome (Mooneyham and Schooler, 2013), while their intrinsic nature constitutes a challenge to their measurement (Smallwood, 2013). Although there is a growing understanding that self-generated states constitute an important feature of human cognition, we lack an agreed upon classification on their defining features (Seli et al., 2018; Christoff et al., 2018), as well as the tools with which to study them (Kucyi, 2018). The current study examined whether it is possible to gain insight into the repertoire of self-generated states by applying advanced machine learning methods to recordings of neural data during periods of wakeful rest, and using these to predict measures of experience.

Our study builds on an emerging literature using advanced analyses techniques to understand the organisation of self-generated cognition. One common method is to reduce the dimensionality of self-reported data in order to identify latent variables that make up participants’ descriptions of their experience. For example, Ruby and colleagues (2013) used principal component analysis (PCA) and identified patterns of self-generated thought that were distinguished by their associations with either the past or the future. Using lag analysis, they demonstrated that these two states had unique correlates with subsequent mood – prospective self-generated thought increased positive mood, while past thoughts were associated with the reverse pattern. Other authors have used a similar approach to establish that prospective self-generated thoughts are often structured and realistic (Stawarczyk et al., 2013) and that they may play a role in the consolidation of personal goals into concrete plans (Medea et al., 2018). Another approach involves the application of clustering techniques to experience sampling data. For example, Andrews-Hanna and colleagues (2013) used hierarchical clustering to determine distinct patterns of negative and self-relevant thought and specific thinking, each with unique psychological correlates. Other studies have employed techniques that explored the temporal features of experience sampling data. For example, Zanesco (2020) employed a temporal clustering method to experience sampling data generated across several different experimental paradigms. They found several patterns of ongoing cognition that were similar across tasks, and identified regularities in the order with which the hidden cognitive states emerge. Our study builds on these findings by applying a temporal decomposition method to neural data recorded during a period of wakeful rest, a situation which has a high prevalence of self-generated mentation (Smallwood et al., 2009) and in which prior studies have successfully linked neural activity to retrospective reports of experience (Gorgolewski et al., 2014; Maillet et al., 2018; Vatansever et al., 2019). Our aim was to determine whether the application of advanced machine learning methods on neural data revealed states that had reliable associations with an individual’s self-reported experience during that period.

In our study, therefore, we measured neural function during wakeful rest in a large cohort of healthy individuals using functional magnetic resonance imaging (fMRI). These participants completed a set of questions related to their experiences during the scan, right at the end of it, while still inside the scanner, and a subset completed questionnaire measures of physical and mental health at a later session. We applied hidden Markov modelling (HMM) to the neural data to reveal distinct brain states corresponding to specific periods of temporally reoccurring patterns of neural mean activity and functional connectivity (Vidaurre et al., 2017a). We then used the dwell-times of the states as predictors in a multivariate multiple regression with the reports given at the end of the scan as the outcome measures. Finally, we investigated how these states are distributed across a multi-dimensional space formed by three well-described neural hierarchies based on the differentiation of whole-brain connectivity patterns (Margulies et al., 2016). The study workflow is presented in Figure 1.

**Figure 1.**
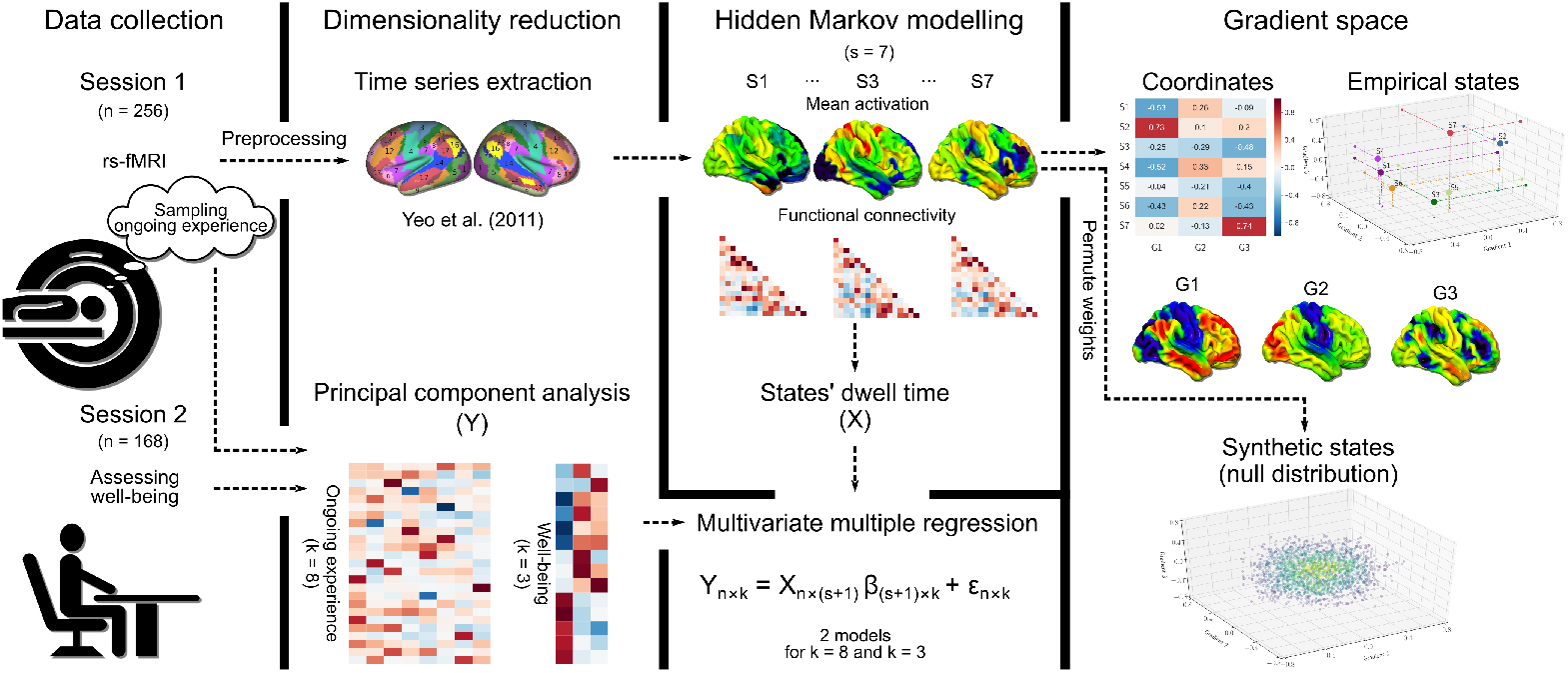
Schematic of the workflow of the current study. We measured neural activity at rest in a sample of n=256 individuals who then completed a retrospective measure of the experiences they had during this period. In a subset of the individuals, we also acquired measures of well-being. We then reduced the dimensionality of both data sets and applied hidden Markov modelling to the neuroimaging time series. We performed a multiple regression analysis in which we calculated the association between the mean dwell-time of each state to the patterns of experiences calculated at rest, as well as to trait measures of well-being. Finally, we calculated the association of the identified brain network states to the first three neurocognitive hierarchies identified by Margulies et al. (2016), as well as to a distribution of synthetic states generated through permutation. k: number of PCA components, s: number of states, G: the three gradients from Margulies et al. (2016)

## Results

### Identifying reoccurring neural states at rest

We applied hidden Markov modelling to the resting state fMRI data (see Methods). This method uses Bayesian inference to identify reoccurring patterns of brain network activity; see Vidaurre et al. (2017b,a) for an empirical validation of this approach. Here, we generated HMM solutions capturing mean activity and functional connectivity patterns within the neural data recorded from our sample. We opted for a 7-state solution, however, our results highlighting associations with behaviour remained broadly similar for a decomposition of 9 states as well. We ran the algorithm 10 times in each case and found that the 7-state solution produced the same decomposition across all iterations and was relatively more stable and had greater split-half reliability than the 9-state solution (see Fig. S2). Accordingly, we focus on the 7-state solution and report a parallel analysis of the 9-state solution in the supplementary Materials (Fig. S5 and S6). Figure 2 shows the spatial description of neural mean activity in each state and the associated cognitive terms produced by a meta-analysis using Neurosynth (Yarkoni et al., 2011), displayed in the form of word clouds.

**Figure 2.**
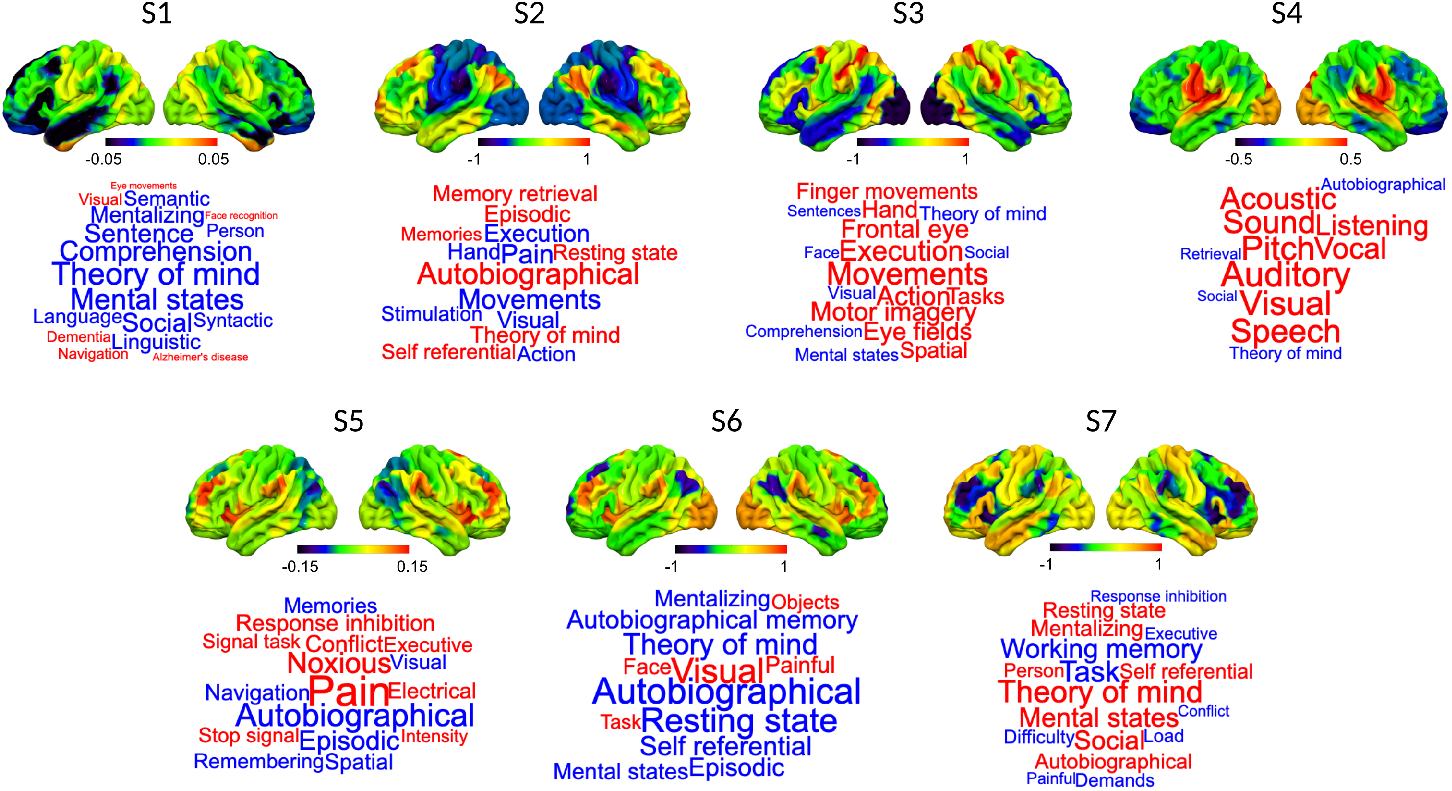
Spatial distribution of mean neural activity characteristic of each state and their meta-analytic associations. The brains show the relative levels of neural activity that are characteristic of each state. Meta-analytic patterns were generated using Neurosynth and are displayed in the form of word clouds. The size of the words reflects the importance of the term, and the colour reflects the direction of the loading (red = positive and blue = negative).

### The relationship between neural states and experience

Having identified a set of relatively stable recurring patterns within our data, we next explored how these naturally occurring neural states are related to the experiences reported by our participants. To answer this question, we performed a multi-variate analysis of co-variance, in which the mean dwell-time of each state was an explanatory variable and the self-reported data was the dependent variable (see Methods for details of the analyses; the specific questions can be found in Table S1). We included motion, age and gender as co-variates of no interest.

This analysis revealed that the mean dwell-time of states 3 and 7 had a significant multivariate association with patterns of reports made by our participants at the end of the scan (F(8, 238) = 2.22, p = 0.027, Wilks’ Λ = 0.931, partial *η*^2^ = 0.069 for state 3 and F(8, 238) = 2.18, p = 0.03, Wilks’ Λ = 0.932, partial *η*^2^ = 0.068 for state 7). These results are presented in Figure 3. It can be seen in the radar plot that state 3 is dominated by task-positive systems (frontal parietal and dorsal attention) as well as the medial temporal subsystem of the default mode network (DMN). Experientially, it was associated with a state of autobiographical planning, reflecting high scores on “Future” and “Problem solving” thoughts. In contrast, state 7 was associated with somatomotor, the core and lateral temporal subsystems of the DMN and the limbic system. This state was associated with reports of negative, intrusive thoughts about the past. Our analysis therefore identified two distinct neural states that were uniquely associated with states associated with experiential foci on different temporal epochs (the past and the future), a pattern that is routinely seen in experience sampling studies (e.g. Ruby et al., 2013; Smallwood et al., 2016; Poerio et al., 2013).

**Figure 3.**
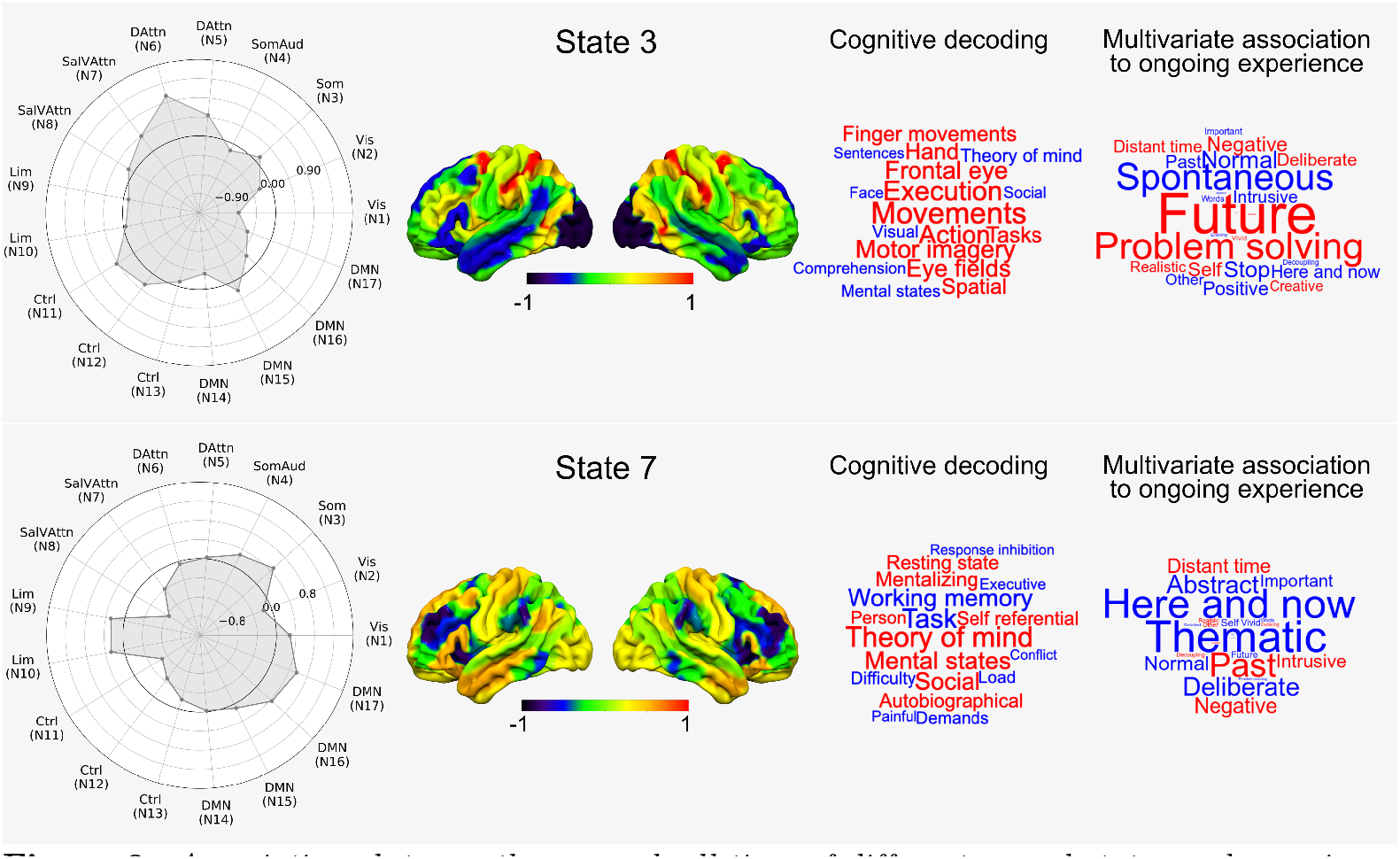
Associations between the mean dwell-time of different neural states and experience. This figure shows the spatial distribution of the two states as well as a meta-analysis of these spatial maps using Neurosynth, which are displayed in the form of word clouds. The word clouds on the right show the pattern of answers associated with each state. In addition, we show, for each state, the relative contributions of each of the networks used in the HMM analyses, in the form of radar plots.

Psychological studies have shown that patterns of ongoing thought often have associations with negative affect, and in particular those associated with the past (Ruby et al., 2013; Smallwood and O’Connor, 2011; Poerio et al., 2013). Based on this evidence, our next analysis aimed at identifying whether the mean dwell-time of the states varied with trait measures of well-being recorded in a subset of these participants. This analysis found that two of the states had a multivariate association with the questionnaire data (state 4, F(3,155) = 2.71, p = 0.047, Wilks’ Λ = 0.95, partial *η*^2^ = 0.05 and state 7, F(3, 155) = 3.74, p = 0.013, Wilks’ Λ = 0.933, partial *η*^2^ = 0.067). These relationships are presented in Figure 4 where it can be seen that state 4 is associated with higher levels of reports of depression and rumination. In contrast, state 7, also identified in our prior analyses, was most strongly linked to trait, state and social anxiety, rumination, depression, as well as greater symptoms linked to autism. Notably, therefore, our analysis identified that state 7, linked to unpleasant thoughts about the past, was also linked to states characterised by psychiatric symptomatology. Our observation of an association between retrospective experiential states and unpleasant affective states is consistent with prior experience sampling studies (Ruby et al., 2013; Smallwood and O’Connor, 2011; Poerio et al., 2013), increasing our confidence in the association between the neural states and patterns of ongoing thought.

Finally, we examined the robustness of our results linking the neural states to the psychological measures using the mean dwell-times of the 9-state solution. In the 9-state solution there were states that were similar to those in the 7-state solution (see Fig. S3). A multivariate analysis showed that state 5, the homologue of state 3 in the 7-state solution, was associated with a similar multivariate pattern of experience, and state 6 (the homologue of state 7 in the 7-state solution) was related to the same pattern of trait measures (Fig. S5). Although no pattern of thoughts was significantly associated with state 6 in the 9-state solution, its experiential correlates were most similar across individuals to those seen in state 7 from the 7-state solution (r = 0.6, p < .001, see Fig. S6). These analyses suggest that associations of neural patterns with both experience and well-being are relatively well-preserved across small changes in the number of states generated through HMM.

**Figure 4.**
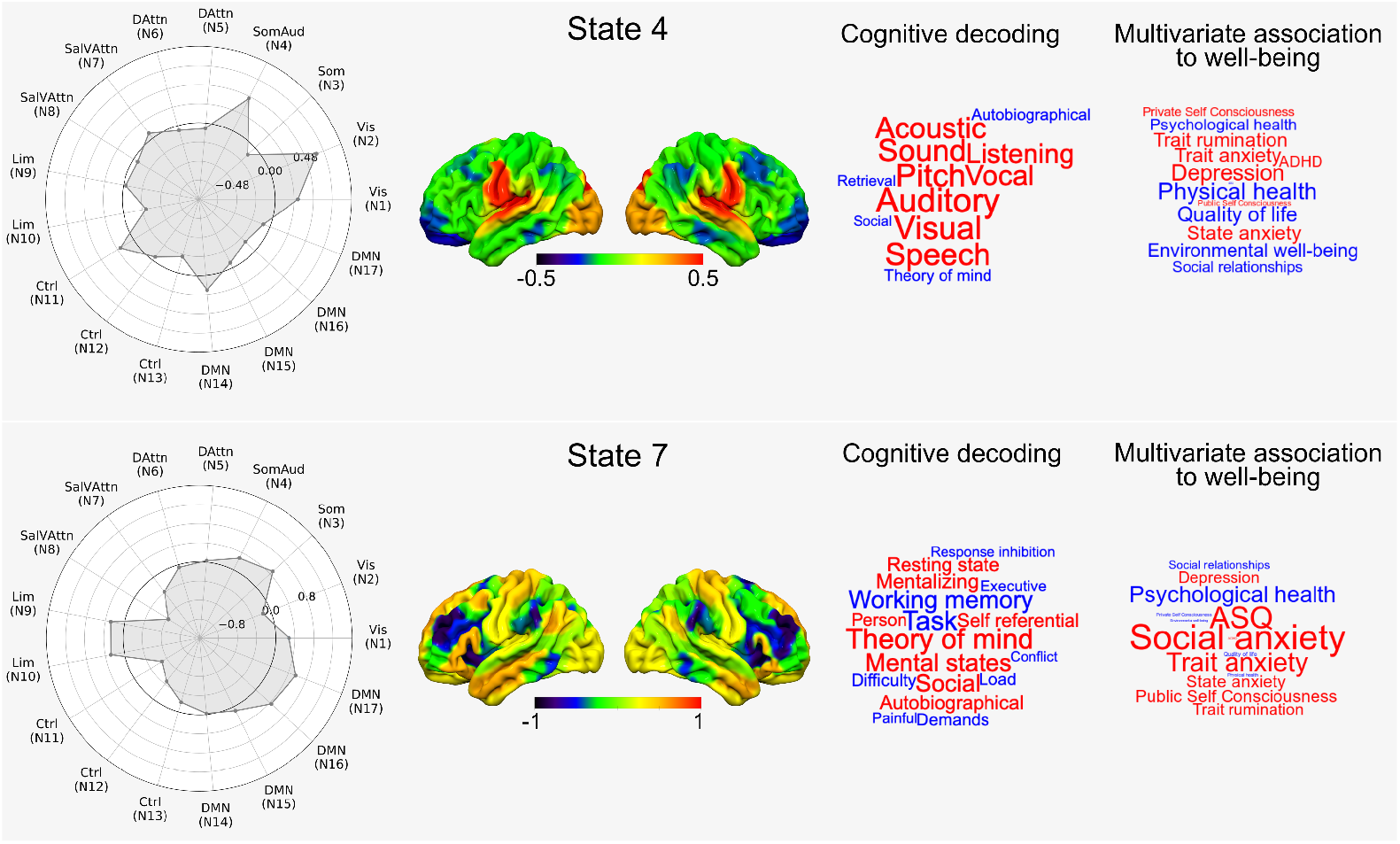
Associations between the mean dwell-time of different neural states and well-being. This figure shows the spatial distribution of the two states as well as a meta-analysis of these spatial maps using Neurosynth, which are displayed in the form of word clouds. The word clouds on the right show the pattern of answers associated with each state. In addition, we show for each state, the relative contributions of each of the networks used in the HMM analyses, in the form of radar plots.

### The relationship between naturally occurring states at rest and neural hierarchies

Having identified network states related to experience and trait measures of well-being, we next explored their association with three well-described neural hierarchies that highlight gradients in connectivity patterns over space. We chose the first three hierarchies from Margulies et al. (2016) which represent the division of unimodal and transmodal systems (gradient 1), the distinction between vision and motor (gradient 2) and the patterns seen during the brain’s response to tasks relative to rest (gradient 3). These were generated using diffusion embedding, a non-linear dimensionality reduction technique, on the Human Connectome Project Data, and identified components describing the maximum variance in functional connectivity patterns (see Methods and initial paper for further details). The spatial distribution of each of these gradients is presented in Figure 5a along with a meta-analysis of these maps using Neurosynth.

**Figure 5.**
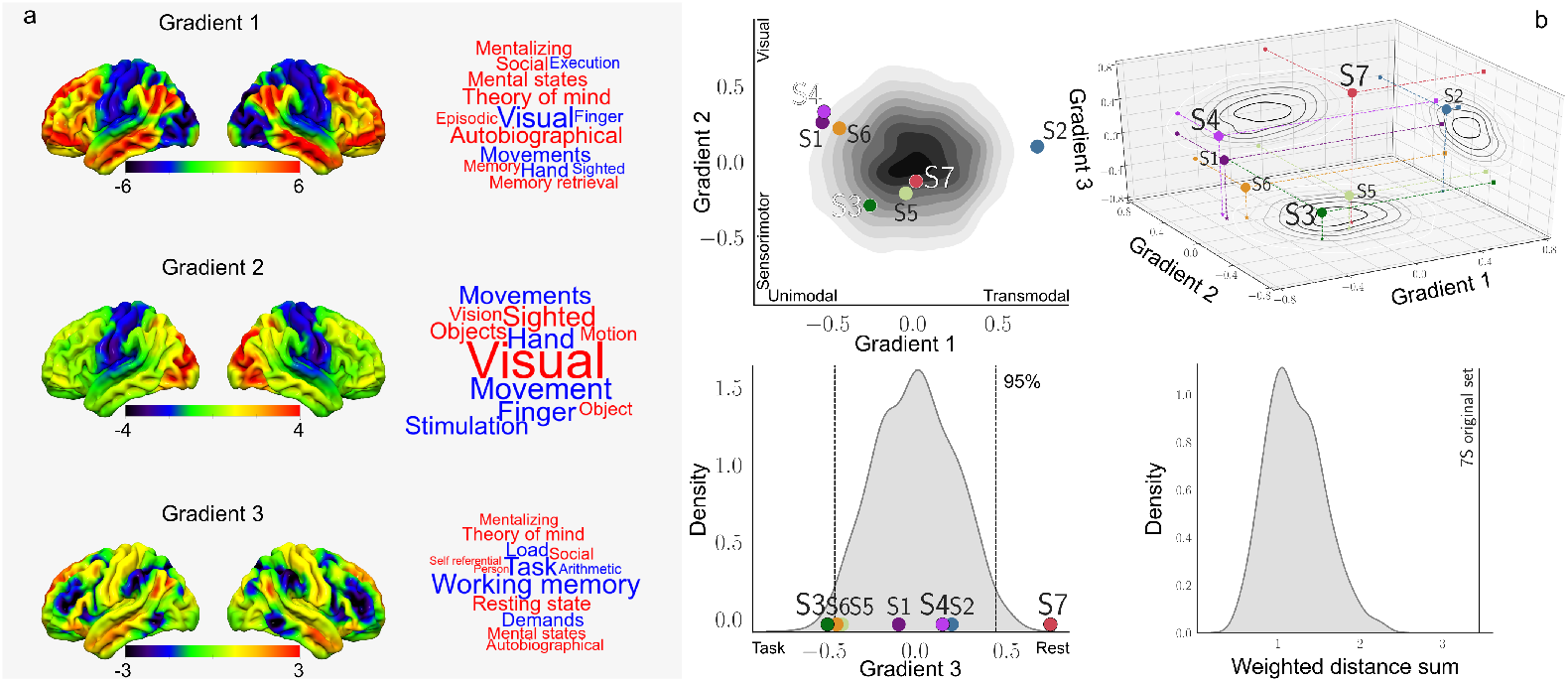
Relationship between naturally occurring states at rest and three well-established cognitive hierarchies. **a**) This panel shows the spatial distribution of the three neurocognitive hierarchies identified by Margulies et al. (2016), as well as a meta-analysis of these spatial maps using Neurosynth, which are displayed in the form of word clouds. Font size represents the strength of the association and font colour its sign (red for positive and blue for negative values). **b**) This panel shows the spatial similarity between the spatial maps describing these neural hierarchies and the set of spatial maps generated from the HMM decomposition of the real data. In these figures, the contour plots projected onto each plane describe the distribution of values generated through permutation. The lower right-hand histogram shows the distribution of synthetic states compared to the real data, in terms of the sum of the weighted distance from the origin of each point in each set. States 3, 4 and 7, whose dwell-time was associated with psychological measures, are highlighted.

The aim of this analysis was to understand whether there is a relationship between the states inferred by the HMM (see Figure 2) and the spatial maps which represent these pre-established hierarchies. To achieve this goal, we calculated the pair-wise similarity between each of the hierarchies and each of the states (see Methods). We compared the associations between the real states inferred in our analyses to a null distribution of synthetic states. The null distribution was generated by randomly permuting the weights describing the relative contribution of each brain region to the mean neural activity captured by each state (see Methods). The results of these analyses are presented on Figure 5b. The contour plot and histogram on the left show where the real data fall on a subset of the dimensions, while the three-dimensional scatter plot show the same data in a combined manner. In these plots, the null distribution generated through permutation is represented as the shaded area in grey.

We also generated a video that shows the data in a 3-d form (https://github.com/tkarapan/hmm-large-scale-hierarchies/blob/master/7S_Null.mp4).

The synthetic states clustered towards the middle of the three dimensions, indicating that states generated in a random manner tend to not be very similar in spatial terms to the three pre-established neurocognitive hierarchies (Margulies et al., 2016). In contrast, the states generated based on real data fall, on average, away from the centre and towards the outer edge of the distribution of synthetic states. Notably, the states with a significant association to experience, states 3 and 7, fall towards the mid-point of both gradient 1 and 2 and, in contrast, are maximally dissociated along the dimension describing how the brain responds to a task; state 3 is more similar to the task positive end of gradient 3, while state 7 shows the reverse. Finally, it can be seen that state 4, which alongside state 7 was associated with trait measures of well-being, falls towards the unimodal end of gradient 1.

To quantify whether the apparent difference between the real and permuted data was robust, we calculated the sum of the weighted distance from the origin for each of the sets of states generated through permutation of this space, and compared this to the same value from the real data. The distribution of these values is presented in the right-hand histogram in Figure 5b, where it can be seen that the weighted distance sum of the real states from the centre of this space is higher than any of the states generated synthetically.

Together, these analyses suggest that states determined through the application of HMM to our real data have an association with the spatial maps describing neural hierarchies that is greater than chance. To identify the robustness of this effect, we performed a parallel analysis based on the 9-state solution to the current data, as well as the published states inferred from applying HMM with the same model parameters on the Human Connectome Project data (Vidaurre et al., 2017b), finding comparable results (see Fig. S4). Together, these analyses show that the states that occur naturally at rest, as approximated through HMM, fall at the extremities of a co-ordinate space defined by well-established neural hierarchies. This result implies that macro-scale hierarchies can be thought of as constraining the state space from which the intrinsic network states emerge.

## Discussion

The current study set out to understand whether it is possible to shed light on the repertoire of self-generated states that an individual engages in, through the application of advanced machine learning methods to neural data recorded during periods of wakeful rest. Our results indicated a significant association between the amount of time an individual spends in two neural states and the experiential reports gained at the end of the scan. One state had features mimicking patterns of neural activity seen during complex tasks (e.g. Duncan, 2010; Power and Petersen, 2013) and was associated with patterns of thoughts focused on future problem solving. A second state highlighted the dominance of the default mode network (Raichle et al., 2001) and was linked to reports of negative intrusive thought. Importantly, this state was also linked to traits associated with negative affect (anxiety, depression and rumination). Together, our study adds to a growing consensus on the utility of methods that explore neural activity from a dynamical perspective, as a tool for understanding patterns of self-generated thought (Kucyi, 2018).

Our analyses, therefore, establish that techniques that parse continuous neural data into time-varying states can be used as a tool to empirically constrain accounts of the qualitative patterns of ongoing thought. By relating the amount of time an individual spent in specific neural states to the pattern of thoughts reported at the end of the scan, we highlighted many of the themes of self-generated thought that are common within the literature. For example, patterns of future problem solving identified in our study are consistent with a prospective bias to ongoing thought that develops in young adulthood (McCormack et al., 2019; Irish et al., 2019) and is common in multiple cultures (Australia, Irish et al., 2019; Belgium, Stawarczyk et al., 2013; China, Song and Wang, 2012; Germany, Ruby et al., 2013; Japan, Iijima and Tanno, 2012; U.K. Smallwood et al., 2009; USA, Baird et al., 2011; Seli et al., 2017). This may reflect a mode of autobiographical planning that is hypothesised to be a potential beneficial outcome of self-generated thought (Baird et al., 2011; Stawarczyk et al., 2013). The second state we identified emphasises negative intrusive thoughts from the distant past and was associated with high levels of trait affective disturbance. This pattern of unhappy rumination may reflect the link between negative affective thought and a focus on the past that is often observed in studies of self-generated thought (Ruby et al., 2013; Smallwood and O’Connor, 2011; Spronken et al., 2016; Poerio et al., 2013). Notably, our study estimated states simply based on neural information without the need to repeatedly sample participants’ experience. In this context, the synergy between the qualitative motifs associated with our neural states and those identified by prior experience sampling studies helps minimise concerns that experience sampling results are in general a consequence of the meta cognitive demands imposed by this technique (Smallwood and Schooler, 2015).

Finally, our study provides a novel view of the relationships between specific macro-scale hierarchies and patterns of spontaneously occurring neurocognitive states. The two states identified that had experiential correlates fell at opposite extremes of a neurocognitive hierarchy that reflects the neural response to tasks. These data suggest that similar neural hierarchies that support general aspects of how individuals respond to the demands of an external task (Duncan, 2010) may continue to do so in the absence of any overt external behaviour. Importantly, this neural hierarchy discriminated between patterns of autobiographical planning, which are thought to be advantageous because they help consolidate personal goals (Medea et al., 2018), and those linked to patterns of rumination, which exacerbate unhappiness (Poerio et al., 2013). Our results thus suggest that the same neural hierarchy that determines the brain’s response to increasing task demands may also discriminate beneficial and detrimental types of self-generated experiences.

Although our study highlights the utility of advanced machine learning methods in the determination of naturally occurring self-generated states, it leaves open several important questions. For example, our application of HMMs allowed us to estimate the occurrence of transient states at rest; however, we only gained a single measure of an individual’s experience. This feature of our experimental design allows us to rule out certain meta-cognitive features of experience sampling as confounding our results (see above); however, it also makes unclear the extent to which the observed states are transient or result from more stable trait-like properties of the individual. In the future, studies could overcome this limitation by sampling an individual’s experience on multiple occasions. This would allow for a more quantified assessment of whether these states occur in a reliable manner, or whether they are subject to more transient influences. Moreover, although our study highlights that neural information can be used to provide a quantified assessment of the repertoire of states that individuals engage in, it leaves open the specific experiential features that these states may include. In our study, we sampled individuals’ thoughts using 25 questions that were refined through a sequence of empirical investigations, yet, it seems likely that these are not an exhaustive list of experiential states, because at least certain features of self-generated experience are probably not captured by the specific items we used. Accordingly, our study establishes that neural information can provide an important window into the types of thoughts individuals may engage in. Nevertheless, future work is needed to establish the most appropriate self-report items to fully appreciate the full range of the ontological features of self-generated experience. Finally, our study leaves open the specific duration that states can take. Our study used fMRI to estimate the neural states and we used a repetition time (TR) that was reasonably slow (3 seconds). This feature of our design acts as a hard limit to the duration of states that our analysis can determine since we would be unable to determine states that lasted substantially shorter durations that that of the TR. In the future, it will be possible to identify states with a shorter duration by performing a similar set of analyses, using a method of neuroimaging such as electro/magnetoencephalography that can acquire neural data in a more rapid manner.

## Methods and Materials

### Participants

277 healthy participants were recruited from the University of York. Informed consent was obtained for all participants and the study was approved by the York Neuroimaging Centre Ethics Committee. 21 participants were excluded from analyses, 1 due to technical issues during the neuroimaging data acquisition and 20 for excessive movement during the fMRI scan (mean framewise displacement (Power et al., 2014) > 0.3 mm and/or more than 15% of their data affected by motion), resulting in a final cohort of n = 256 (169 females, *μ*_*age*_ = 20.7 years, *σ*_*age*_ = 2.4).

### Behavioural methods

We sampled participants’ experience during the resting state fMRI scan by asking them to retrospectively report their thoughts at the end of the scan. Experience was measured using a 4-scale Likert scale, with the question order randomised (all 25 questions are shown in Table S1). In a subset (n=168) of the final cohort, we also assessed their physical and mental health by administering well-established questionnaire measures at a later separate session outside of the scanner. Details about each questionnaire are presented in SI.

### MRI data acquisition

MRI data were acquired on a GE 3 Tesla Signa Excite HDxMRI scanner, equipped with an eight-channel phased array head coil at York Neuroimaging Centre, University of York. For each participant, we acquired a sagittal isotropic 3D fast spoiled gradient-recalled echo T1-weighted structural scan (TR = 7.8 ms, TE = minimum full, flip angle = 20°, matrix = 256×256, voxel size = 1.13×1.13×1 mm^3^, FOV = 289×289 mm^2^). Resting-state functional MRI data based on blood oxygen level-dependent contrast images with fat saturation were acquired using a gradient single-shot echo-planar imaging sequence with the following parameters; TE = minimum full (≈19 ms), flip angle = 90°, matrix = 64×64, FOV = 192×192 mm^2^, voxel size = 3×3×3 mm^3^, TR = 3000 ms, 60 axial slices with no gap and slice thickness of 3 mm. Scan duration was 9 minutes, which allowed us to collect 180 whole-brain volumes.

### fMRI data pre-processing

Functional MRI data pre-processing was performed using SPM12 (http://www.fil.ion.ucl.ac.uk/spm) and the CONN toolbox (v.18b) (https://www.nitrc.org/projects/conn) (Whitfield-Gabrieli and Nieto-Castanon, 2012) implemented in Matlab (R2018a) (https://uk.mathworks.com/products/matlab). Pre-processing steps followed CONN’s default pipeline and included motion estimation and correction by volume realignment using a six-parameter rigid body transformation, slice-time correction, and simultaneous grey matter (GM), white matter (WM) and cerebrospinal fluid (CSF) segmentation and normalisation to MNI152 stereotactic space (2 mm isotropic) of both functional and structural data. Following pre-processing, the following potential confounding effects were removed from the BOLD signal using linear regression: 6 motion parameters calculated at the previous step and their 1st and 2nd order derivatives, volumes with excessive movement (motion greater than 0.5 mm and global signal changes larger than z = 3), signal linear trend, and five principal components of the signal from WM and CSF (CompCor approach, Behzadi et al., 2007). Finally, data were band-pass filtered between 0.01 and 0.1 Hz. No global signal regression was performed.

### Dimensionality reduction

In order to reduce the dimensionality structure of the neuroimaging data, we used the 17 functional network parcellation from Yeo et al. (2011), slightly eroded the parcels to avoid signal leakage from neighbouring parcels, masked them with subject specific grey matter masks, and calculated the average signal from the unsmoothed data within each parcel for each volume per participant. The acquired time series were then standardised and concatenated to form a “(256 participants × 180 volumes) × 17 parcels” matrix that was used as input to the HMM algorithm.

For the behavioural data, we applied a principal component analysis (PCA) with varimax rotation to the scores describing the participants’ experience at the end of the resting state scan. This revealed eight components with eigenvalues greater than 1. We followed the same procedure for the trait measures of well-being that we acquired for a sub-sample of our cohort, which identified three principal components in their structure (see Figure S1 for the loadings of both decompositions in the form of word clouds). The component scores were used as dependent variables in two separate multivariate multiple regressions, run at a later stage in order to investigate the relationship between ongoing experience and well-being respectively, with the mean dwell-times of the dynamic neural states as described by our HMM decomposition.

### Hidden Markov model

To characterise the dynamics of neural activity, we applied hidden Markov modelling to the concatenated time series of the 17 parcels (Yeo et al., 2011). The inference of the model parameters was based on variational Bayes and the minimisation of free energy, as implemented in the HMM-MAR toolbox (Vidaurre et al., 2016). The HMM’s inference assigns state probabilities to each time point of the time series (i.e. reflecting how likely each time point is to be explained by each state) and estimates the parameters of the states, where each state has its own model of the observed data. Each state was represented as a multivariate Gaussian distribution (Vidaurre et al., 2017a), described by its mean and covariance. Inference was run at the group level, such that the state spatial descriptions are defined across subjects, whereas the temporal activation of the states is defined at the subject level. This allowed us to discover dynamic temporal patterns of whole-brain activity and functional interactions along with their occurrence (state time series) and transition probabilities for the duration of the whole resting state fMRI scan. Detailed information about the HMM implementation and the variational Bayes inference can be found in (Vidaurre et al., 2016, 2017b,a). As with other probabilistic unsupervised learning methods (e.g. independent component analysis), HMM is sensitive to initial values. In order to account for HMM run-to-run variability, we ran the algorithm 10 times and selected the iteration with the lowest free-energy at the end of the inference. The summary statistic describing the states’ dwell-times was computed after hard classifying the states as being active or not by using Viterbi decoding. The mean dwell-time of each state was then used as an explanatory variable in subsequent multivariate generalised linear models. Dwell-times were standardised and any values > 2.5*σ* were substituted with the mean (*μ* = 0) to minimise the effect of potential outliers. All analyses controlled for age, gender and motion during the resting state fMRI scan.

### Projection of state maps to a 3-dimensional space of neural hierarchies

Following the HMM inference, each state S can be characterised by parameters (*μ*_*s*_, Σ_*s*_), where *μ*_*s*_ is a vector containing the activation/weights (with respect to the average) of each of the 17 parcels used in our analyses and Σ_*s*_ is the covariance matrix describing their functional interactions. Using the weights of each parcel, we obtained each state’s “mean activity” spatial map (Fig. 2) and calculated its spatial similarity to the first three gradient maps from Margulies et al. (2016). These maps were produced by a diffusion embedding algorithm, applied on data from the Human Connectome Project, they are orthogonal, and highlight topographies of information exchange based on the differentiation of connectivity patterns. Similarity was calculated as the pairwise correlation between each state spatial map and gradient and was used as the state’s co-ordinate for the corresponding dimension in gradient space. Aiming to construct a summary metric describing the topography of the states in this space, we Fisher-z transformed the correlation values (used as gradient space co-ordinates), calculated the distance of each state from the origin weighted by its maximum co-ordinate (as a way to differentiate between points lying on the surface of the same sphere), and added the weighted distances to produce a sum for the set of all states

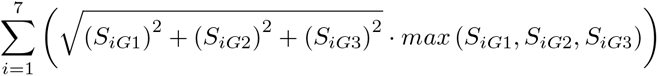

where *S*_*iG*1_, *S*_*iG*2_, *S*_*iG*3_ are the Fisher’s z correlation values between the spatial maps of state *i* and gradient 1, 2, and 3 respectively.

### Null distribution

In order to compute a null distribution, we randomly permuted the parcel weights, constructed the synthetic states’ spatial maps and projected them into gradient space, by calculating their similarity to the gradients in the same way as before. We ran 300 permutations, producing 2100 synthetic states and calculated the weighted distance sum of each set in a similar manner as with the set of empirical states.

## Code availability

Code used for the HMM analyses can be accessed on https://github.com/OHBA-analysis/HMM-MAR.

## Data availability

The data that support the findings of this study are available from the corresponding author upon reasonable request.

## Acknowledgments

JS was supported by European Research Council (WANDERINGMINDS - 646927). The authors would like to thank Mladen Sormaz, Charlotte Murphy and Hao-Ting Wang for their contribution to data acquisition.

