## Supplementary Material for "The psychological correlates of distinct neural states occurring during wakeful rest"

### Emergence of neural dynamics within a co-ordinate space of large-scale neural hierarchies

#### This PDF file includes:

Supplementary text
Figures S1 to S6
Table S1
References for SI citations

### Supporting Information Text

### SI Materials and Methods

**Experience sampling questions.** We asked participants to answer 25 questions shown in Table S1 at the end of the resting state functional Magnetic Resonance Imaging (rs-fMRI) scan, relating to their thoughts during this period. Answers were given on a 4-scale Likert scale ranging from "Not at all" to "Completely".

**Physical and mental health questionnaires.** Quality of life, physical and psychological health, social relationships and environmental well-being were measured by the World Health Organization Quality of Life WHOQOL-BREF instrument Organization et al. (2012). Private and public self-consciousness and social anxiety were assessed using the Self-Consciousness scale Scheier and Carver (2013), state and trait anxiety by the State-Trait Anxiety inventory Spielberger and Gorsuch (1983) and trait rumination by the Ruminative response scale Treynor et al. (2003). Finally, depression was measured using the CES-D scale Radloff (1977), autism by the Autism Spectrum Quotient Baron-Cohen et al. (2001) and ADHD by the World Health Organization Adult ADHD Self-Report scale Kessler et al. (2005).

### SI Results

**HMM stability.** To evaluate the stability of our HMM decompositions, we ran the algorithm 10 times and investigated the similarity of the models. Similarity was measured as the correlation between the state time series after optimally reordering the states of each model using the Munkres assignment algorithm Munkres (1957). We found that the 7-state solution produced the same decomposition across all 10 iterations. We repeated the procedure for number of states s = 9 and found that the stability of the solutions was reduced (average similarity $\mu_9 = 0.80$). We then divided our sample randomly into two halves and found that the 7-state solutions were again relatively more reliable (7-states: $\mu_{7split1} = 0.90$ and $\mu_{7split2} = 0.94$, 9-states: $\mu_{9split1} = 0.71$ and $\mu_{9split2} = 0.80$, Fig. S2).

**9-state HMM dwell-times and relation to behaviour.** We performed the same analyses aimed at identifying how the mean dwell-time of the dynamic states varied with measures of ongoing experience and trait measures of well-being for an HMM decomposition of 9 states. These analyses revealed that the dwell-time of states 1 and 5 was associated with differential patterns of reports made by our participants at the end of the scan ($F(8,236) = 2.14$, $p = 0.033$, Wilks' $\Lambda = 0.933$, partial $\eta^2 = 0.067$ for state 1 and $F(8,236) = 2.1$, $p = 0.037$, Wilks' $\Lambda = 0.934$, partial $\eta^2 = 0.066$ for state 5 accordingly). We also found that three of the states had a multivariate association with trait measures of well-being (state 1, $F(3,153) = 3.16$, $p = 0.027$, Wilks' $\Lambda = 0.942$, partial $\eta^2 = 0.058$, state 6, $F(3,153) = 4.66$, $p = 0.004$, Wilks' $\Lambda = 0.916$, partial $\eta^2 = 0.084$, and state 8, $F(3,153) = 2.77$, $p = 0.044$, Wilks' $\Lambda = 0.948$, partial $\eta^2 = 0.052$). These results are presented in Figure S5.

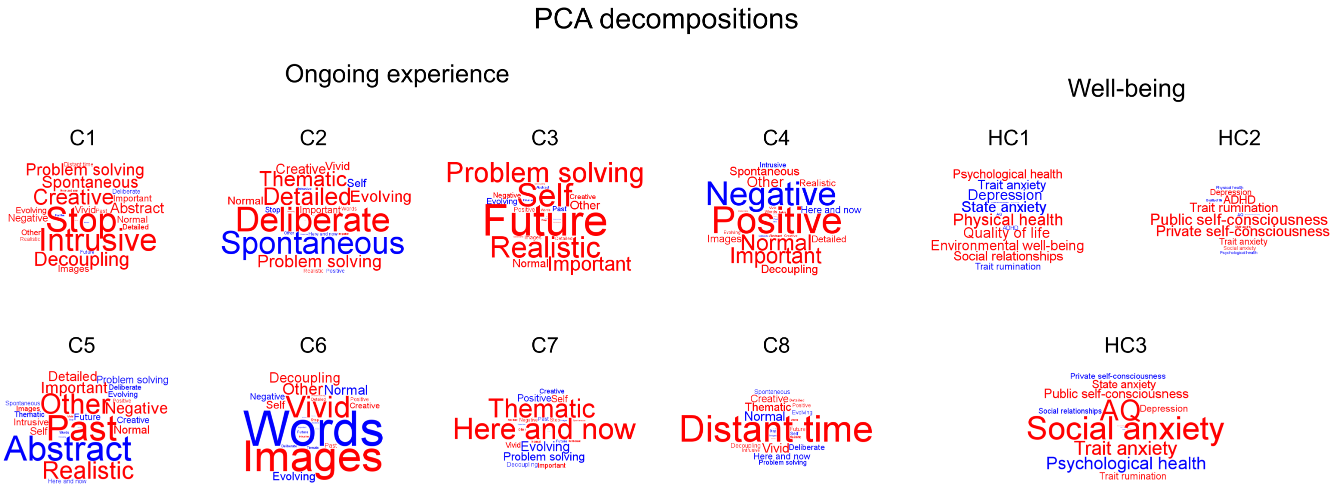


**Figure S1.** *Behavioural variables.* Component weights from principal component analyses on the scores describing the participants' experience at the end of the resting state scan and trait measures of well-being, visualised using a word cloud. Font size represents the strength of the association and font colour its sign (red for positive and blue for negative values).

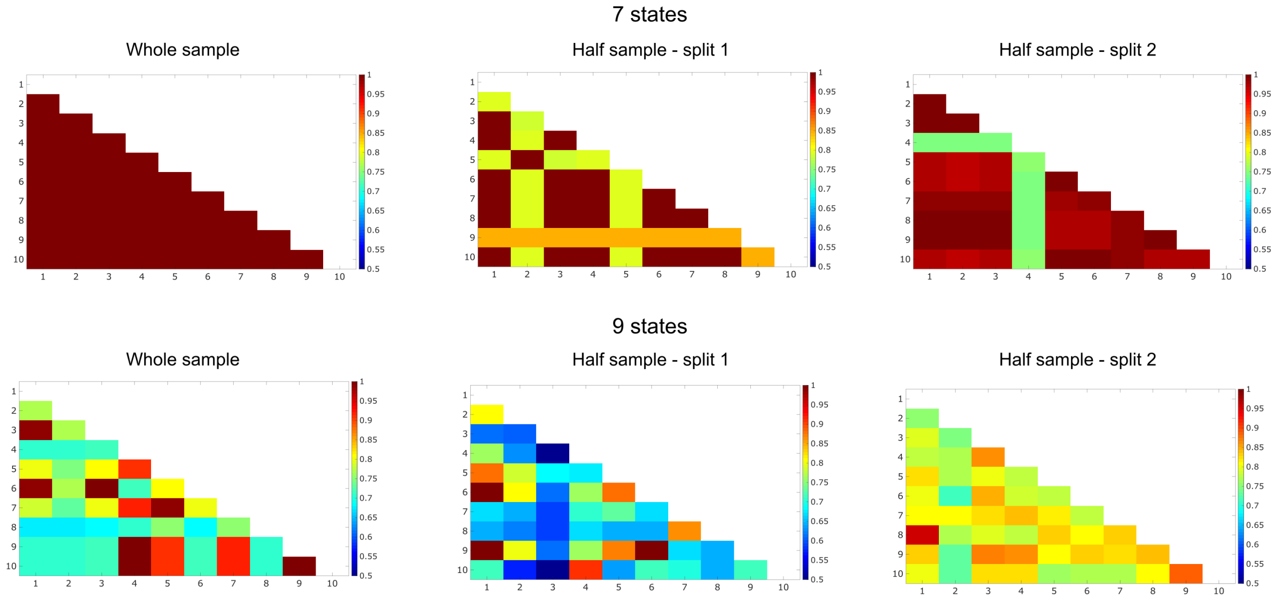


**Figure S2.** *Hidden Markov models similarities*. Similarities between the hidden Markov models after multiple runs of the algorithm for 7 and 9 states using the whole sample and split-halves.

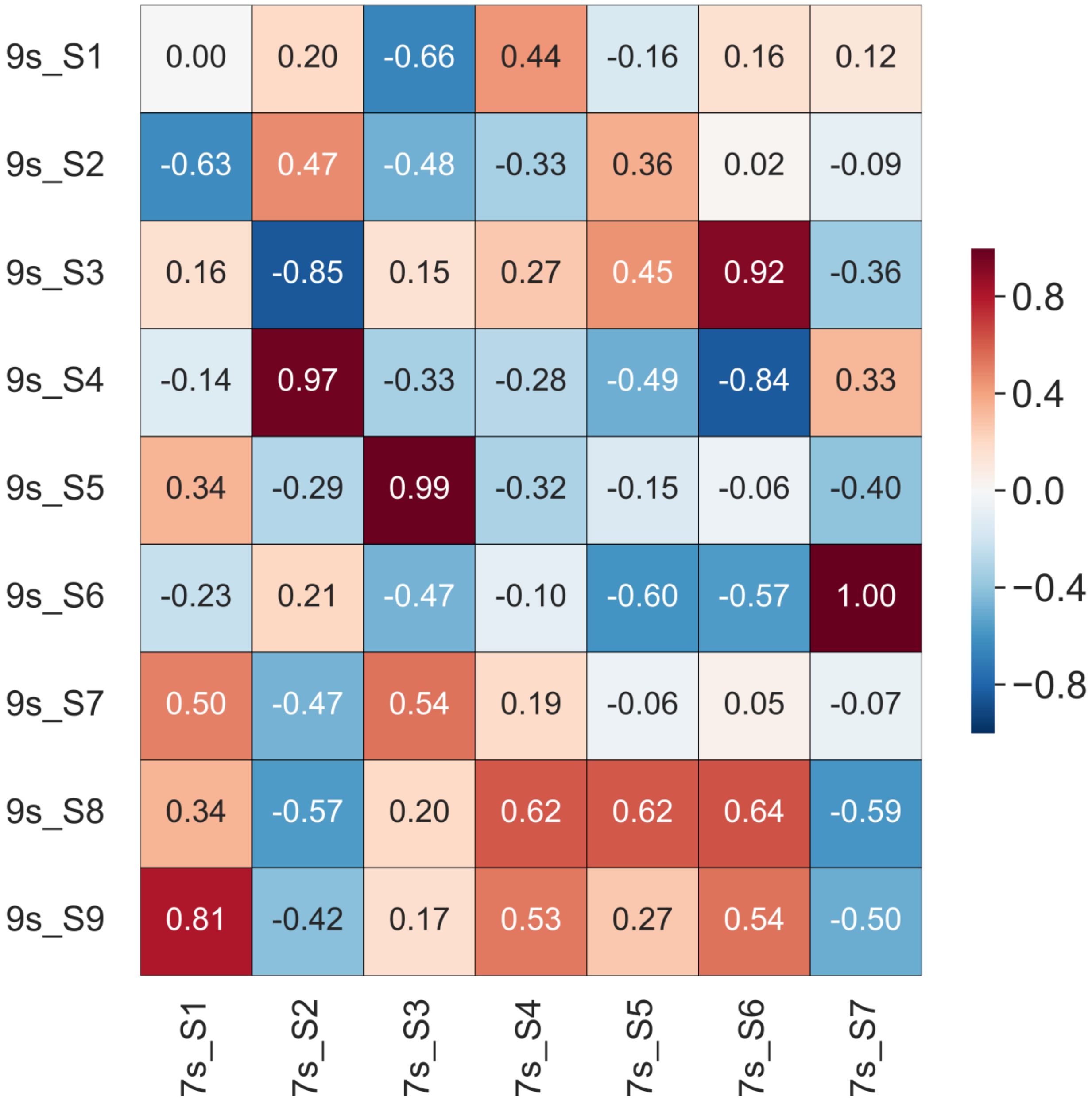


**Figure S3.** *Hidden Markov model states' spatial maps similarities*. Pair-wise correlations between the state spatial maps from an HMM decomposition of our data for 7 and 9 states.

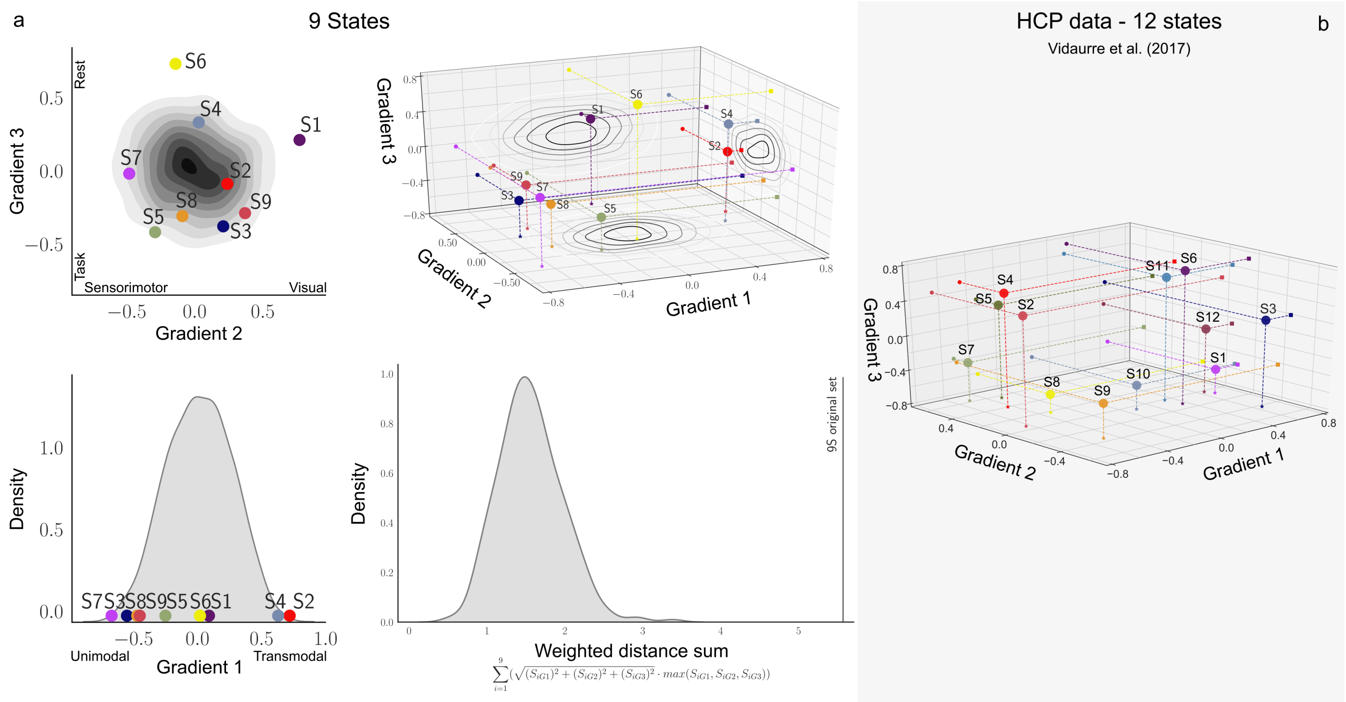


**Figure S4.** *Relationship between naturally occurring states at rest and three well-established cognitive hierarchies, for different HMM decompositions*. **a**) This panel shows the spatial similarity between the spatial maps describing three large-scale neurocognitive hierarchies Margulies et al. (2016) and the set of spatial maps generated from a 9-states HMM decomposition of the real data. In these figures the contour plots projected onto each plane describe the distribution of values generated through permutation. The lower right-hand histogram shows the topology for each set of states generated synthetically as well as for the real data, represented as the sum of the weighted distance from the origin of each point in each set. **b**) This panel shows a three-dimensional scatter plot of the similarities between the functional gradient spatial maps and 12 states inferred from running an HMM on the Human Connectome Project data Vidaurre et al. (2017).

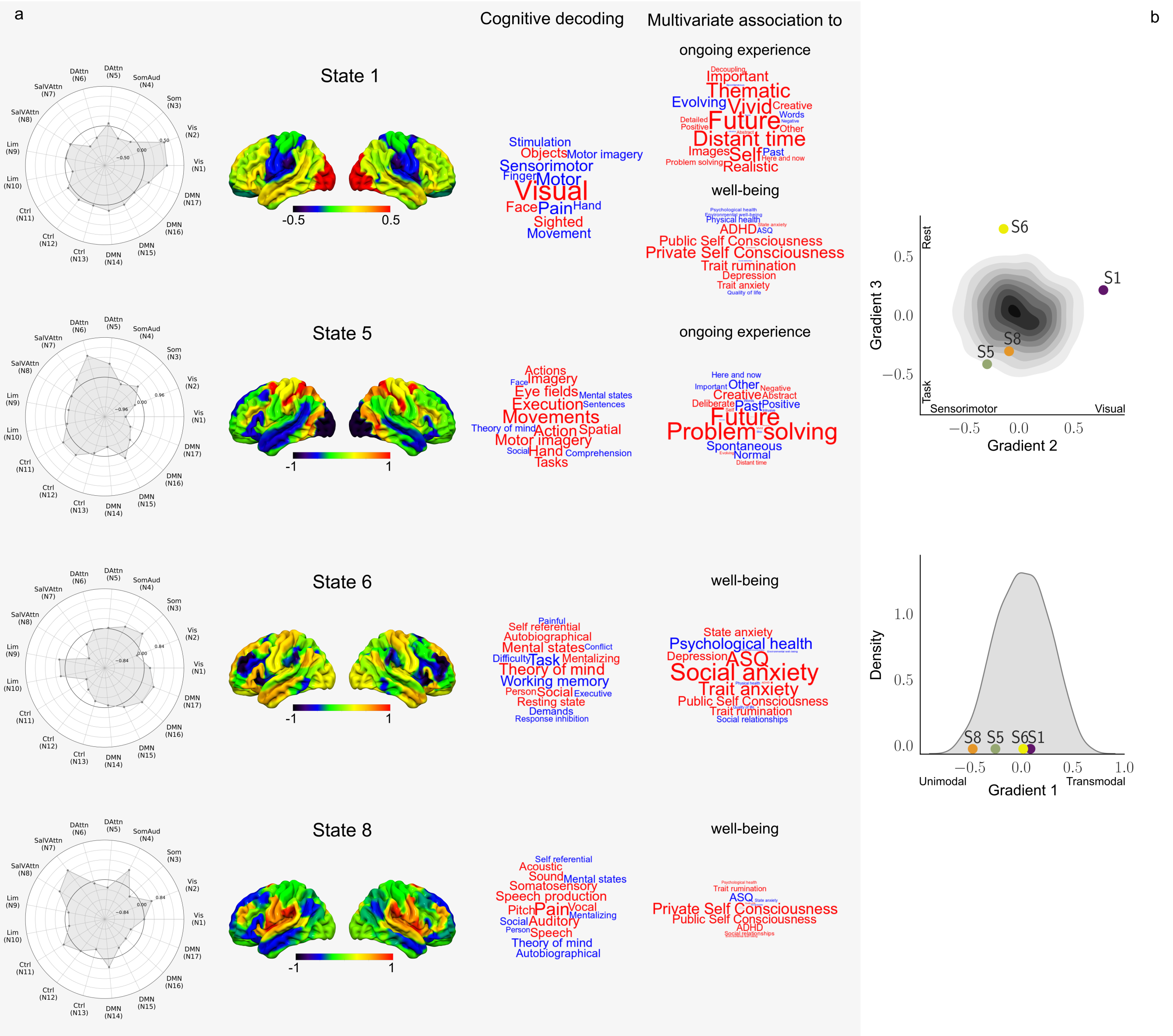


**Figure S5.** *Associations between the mean dwell-time of neural states with experience and well-being, and their relationship to neurocognitive hierarchies, for the 9-states HMM decomposition.* **a**) This panel shows the spatial distribution of the states as well as a meta-analysis of these spatial maps using Neurosynth which is displayed in the form of word clouds. The word clouds on the right show the pattern of answers associated with each state. Font size represents the strength of the association and font colour its sign (red for positive and blue for negative values). In addition, we show for each state, the relative contributions of each of the networks used in the HMM analyses, in the form of radar plots. **b**) This panel shows the spatial similarity between the spatial maps describing these neural hierarchies and the spatial maps generated from the real data.

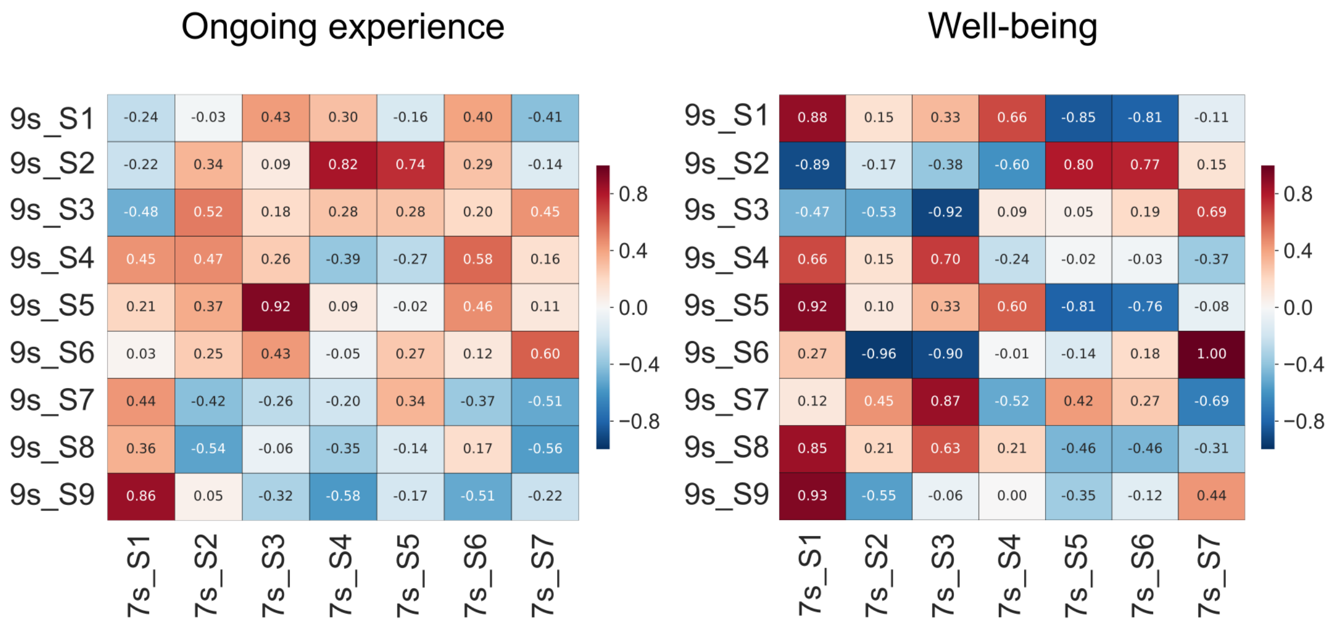


**Figure S6.** *Similarities between the 7-state and 9-state dwell-time experiential correlates*. Heat maps showing the correlations between the multivariate patterns of experience and well-being associated with the states' mean dwell-time from the 7-state and 9-state solutions.

**Table S1. Experience sampling questions asked at the end of the resting state fMRI scan.**

| Dimension | Question<br>(My thoughts) |
|---|---|
| Vivid | ... were vivid as if I was there |
| Normal | ... were similar to thoughts I often have |
| Future | ... involved future events |
| Negative | ... were about something negative |
| Detail | ... were detailed and specific |
| Words | ... were in the form of words |
| Evolving | ... tended to evolve in a series of steps |
| Spontaneous | ... were spontaneous |
| Positive | ... were about something positive |
| Images | ... were in the form of images |
| Other | ... involved other people |
| Past | ... involved past events |
| Deliberate | ... were deliberate |
| Self | ... involved myself |
| Stop | ... were hard for me to stop |
| Distant time | ... were related to a more distant time |
| Abstract | ... were about ideas rather than events or objects |
| Decoupling | ... dragged my attention away from the external world |
| Important | ... were on topics that I care about |
| Intrusive | ... were intrusive |
| Problem solving | ... were about solutions to problems (or goals) |
| Here and now | ... were related to the here and now |
| Creative | ... gave me a new insight into something I have thought about before |
| Realistic | ... were about an event that has happened or could take place |
| Theme | ... at different points in time were all on the same theme |
